# A molecular toolbox to study progesterone receptor signaling

**DOI:** 10.1101/2023.07.20.549847

**Authors:** Marleen T. Aarts, Muriel Wagner, Tanne van der Wal, Antonius L. van Boxtel, Renée van Amerongen

## Abstract

Progesterone receptor (PR) signaling is required for mammary gland development and homeostasis. A major bottleneck in studying PR signaling is the lack of sensitive assays to measure and visualize PR pathway activity both quantitatively and spatially. Here, we develop new tools to study PR signaling in human breast epithelial cells. First, we generate optimized Progesterone Responsive Element (PRE)-luciferase constructs and demonstrate that these new reporters are a powerful tool to quantify PR signaling activity across a wide range of progesterone concentrations in two luminal breast cancer cell lines, MCF7 and T47D. We also describe a fluorescent lentiviral PRE-GFP reporter as a novel tool to visualize PR signaling at the single-cell level. Our reporter constructs are sensitive to physiological levels of progesterone. Second, we show that low background signaling, and high levels of PR expression are a prerequisite for robustly measuring PR signaling. Increasing PR expression by transient transfection, stable overexpression in MCF7 or clonal selection in T47D, drastically improves both the dynamic range of luciferase reporter assays, and the induction of endogenous PR target genes as measured by qRT-PCR. We find that the PR signaling response differs per cell line, target gene and hormone concentration used. Taken together, our tools allow a more rationally designed approach for measuring PR signaling in breast epithelial cells.

## Introduction

The ovarian hormones estrogen and progesterone are essential for the dynamic regulation of mammary gland development and function [1–4]. Although estrogen signaling has received most attention, recent discoveries suggest that progesterone signaling plays a previously underappreciated role in breast cancer [2,5]. The progesterone signaling cascade is active in hormone-sensitive luminal cells, where it regulates a number of critical cellular processes that include the activation of paracrine signaling pathways to induce cell proliferation [6,7]. How exactly progesterone regulates these processes is still incompletely understood.

The progesterone receptor (PR) is a member of the nuclear hormone receptor subfamily that also includes the estrogen receptor (ER), mineralocorticoid receptor (MR), glucocorticoid receptor (GR) and androgen receptor (AR) [8]. The natural PR ligand, progesterone (P4), is a systemic hormone that functions at concentrations ranging from ∼50-350pM during menopause up to 1μM during pregnancy [2,9]. As with all nuclear hormone receptors, progesterone signals by binding to its receptor, PR, which leads to dimerization and nuclear translocation of the hormone-bound receptor. Once in the nucleus, PR binds to specific progesterone responsive DNA elements (PREs) as part of a larger transcription factor complex to regulate a PR-responsive transcriptional program [6,9].

Hormone receptor positive luminal breast cells express two PR isoforms, PR-A and PR-B, which are both encoded by a single *PGR* gene [10,11]. Being transcribed from the proximal start codon, PR-B is 933 amino acids in length. PR-A is transcribed from an alternative, more distal, start codon and consequently lacks the N-terminal 164 amino acids. As a result, both isoforms contain a progesterone-binding pocket and a DNA binding domain, but only PR-B contains the transactivation domain that drives PR-dependent gene regulation [12]. Together, PR-A and PR-B are responsible for overall PR signaling activity [11].

Despite the fact that progesterone was discovered more than 90 years ago and PR was first described over 50 years ago, it has been a challenge to dissect the molecular mechanisms of PR signaling and PR-driven transcriptional activation in the breast epithelium [7,13–16]. This will be essential, however, in order to understand the role of PR in in mammary gland biology during health and disease.

Several factors currently hamper progress in studying PR signaling. A major hurdle that has yet to be taken is the generation of non-transformed, PR-positive breast cell lines, as primary breast epithelial cells typically lose hormone receptor expression when cultured *in vitro* [17]. Primary 3D cultures of normal human breast organoids grown in matrigel that were reported to contain cells with PR expression were unable to induce expression of known PR target genes, including *WNT4* and *RANKL* [7]. Fresh *ex vivo* culture of primary human breast tissue fragments did reveal a preserved hormone response [15], but this culture system is transient, not easily scaled and incompatible with longer term experiments. As a result, most insights into PR signaling have been acquired using the PR-positive luminal breast cancer cell lines MCF7 and T47D. Even these studies suffer from a scarcity of optimized, sensitive, and specific molecular tools to study PR signaling. For instance, inconsistent progesterone induced transcriptional responses measured by qRT-PCR have been reported [15,17], and the few available PRE luciferase reporter constructs show low inducibility and high response variation [18–21]. Furthermore, most *in vitro* studies have used (synthetic) progesterone concentrations in the nM range to study the molecular mechanisms of PR signaling. The question therefore arises if the experimentally measured effects in these studies reflect responses to more physiological progesterone concentrations.

Here, using MCF7 and T47D cell lines, we test and optimize existing approaches to measure PR signaling. We also develop new bioluminescent and fluorescent reporter tools to measure PR signaling activity robustly and quantitatively in either a population based setting or with single cell resolution. We describe optimized conditions that allow robust detection of endogenous PR target gene induction. Taken together, this work serves as a first step to open up new experimental opportunities for the breast (cancer) research field that will allow outstanding questions about PR signaling to be addressed in the near future.

## Results

### PR localization is an imperfect indicator of PR signaling activity

According to the textbook, nuclear hormone receptors – including PR – are predominantly located in the cytoplasm in the absence of ligand, while dimerization and nuclear translocation occurs upon hormone binding. Although this dogma has been challenged [8], immunostaining for PR is frequently used to detect PR expression in healthy mammary epithelial or breast cancer cells [6,22,23]. We reasoned that if PR relocalization indeed occurs in response to progesterone stimulation, quantifying the changes in nuclear PR intensity by immunofluorescence staining should be a useful measure of PR signaling activity.

To test this assumption, we visualized endogenous PR protein in MCF7 and T47D breast cancer cells in response to treatment with the synthetic progestin and PR agonist R5020 (Fig1). In contrast to previous reports [24,25], we found that PR was already predominantly located in the nucleus in unstimulated conditions, although we observed a large variance in nuclear intensity between individual cells in both MCF7 as well as T47D cells (Fig1a – d). Qualitatively, R5020 treatment led to an increase in perceived staining intensity, but quantitatively we only measured an average 1.20-fold (MCF7) and 1.32-fold (T47D) increase compared to unstimulated cells, while variance remained in the same range (Fig1b,d). Treating the cells with R5020 together with the PR antagonist RU486 did not prevent PR from translocating to the nucleus (mean nuclear PR intensity 1.27-fold increase compared to control in both MCF7 and T47D cells) (Fig1b,d). From this, we conclude that nuclear accumulation of PR is a suboptimal readout for quantitative measurements of PR signaling activity. To gain more insight into PR signaling levels and dynamics – including its downstream effects, we therefore decided to develop more robust tools to quantitatively measure the activity and strength of the PR signaling response.

**Fig.1.**
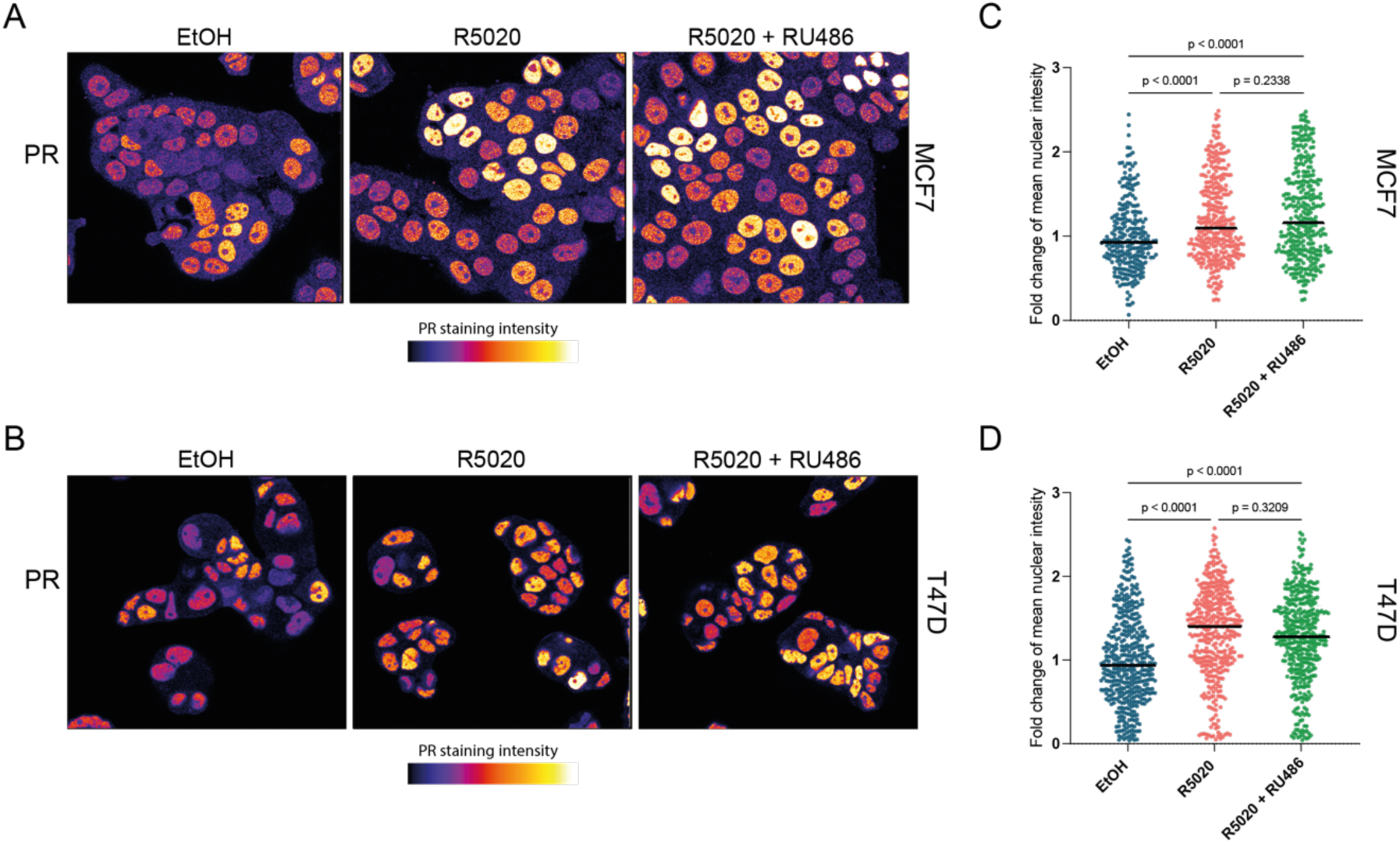
Quantitative measurements of nuclear PR abundance. a,b) Representative confocal microscopy images showing endogenous PR expression detected by immunofluorescence staining of MCF7 (a) or T47D (b) cells treated with EtOH (control), 20nM R5020, or 20nM R5020 and 100nM RU486 for 2 hours. c-d) Quantification of the mean nuclear intensity of the PR staining in MCF7 (c) or T47D (d) cells treated as in A and B (nuclei were segmented using the DAPI signal (not shown)). Each datapoint represents a single nucleus, black bars represent the mean intensity (250-400 cells per condition from N=2 independent experiments).

### PR expression levels determine the strength of the PR signaling response

Generally speaking, (luciferase) reporter gene assays offer a powerful approach for measuring signaling activity at the transcriptional level. This method has also been used for PR signaling, but the inducibility of existing PRE luciferase constructs is relatively low and varies considerably between cell lines [18–21]. Therefore, we set out to optimize a dual luciferase reporter assay to quantify PR signaling activity in MCF7 and T47D cells.

We started with a previously generated construct that contains two consensus PR binding sites (2xPRE) upstream of a minimal thymidine kinase (TK) promoter (Fig2a) [21,26]. Transient transfection of this reporter into MCF7 cells did not significantly induce reporter signal in response to treatment with 20nM R5020 (Fig2a). Including two additional PR binding sites (4xPRE) did little to improve detection (Fig2a,b). Since endogenous PR levels are relatively low in MCF7 cells (SupFig1) [28], we hypothesized that increasing PR expression might result in a stronger response. Indeed, co-transfection of a PR expression plasmid improved the dynamic range of both the 2xPRE and the 4xPRE reporter, allowing us to measure a clear and statistically significant induction of over 100-fold in response to R5020 treatment (Fig2b +PR). Induction was PR specific, since treatment with both R5020 and RU486 abolished the luciferase signal in both -PR and +PR conditions (Fig2b). Therefore, PR co-transfection was used for all further experiments with wildtype MCF7 cells unless noted otherwise. Of note, PR co-transfection did not improve the dynamic range of our PRE luciferase reporters in T47D cells (Fig2c). Even without PR co-transfection T47D cells already showed a clear and statistically significant induction up to 100-fold (4xPRE, Fig2c). This is in line with the fact that endogenous *PGR* expression in T47D cells is more than 8-fold higher than in MCF7 cells (SupFig1).

**Fig.2.**
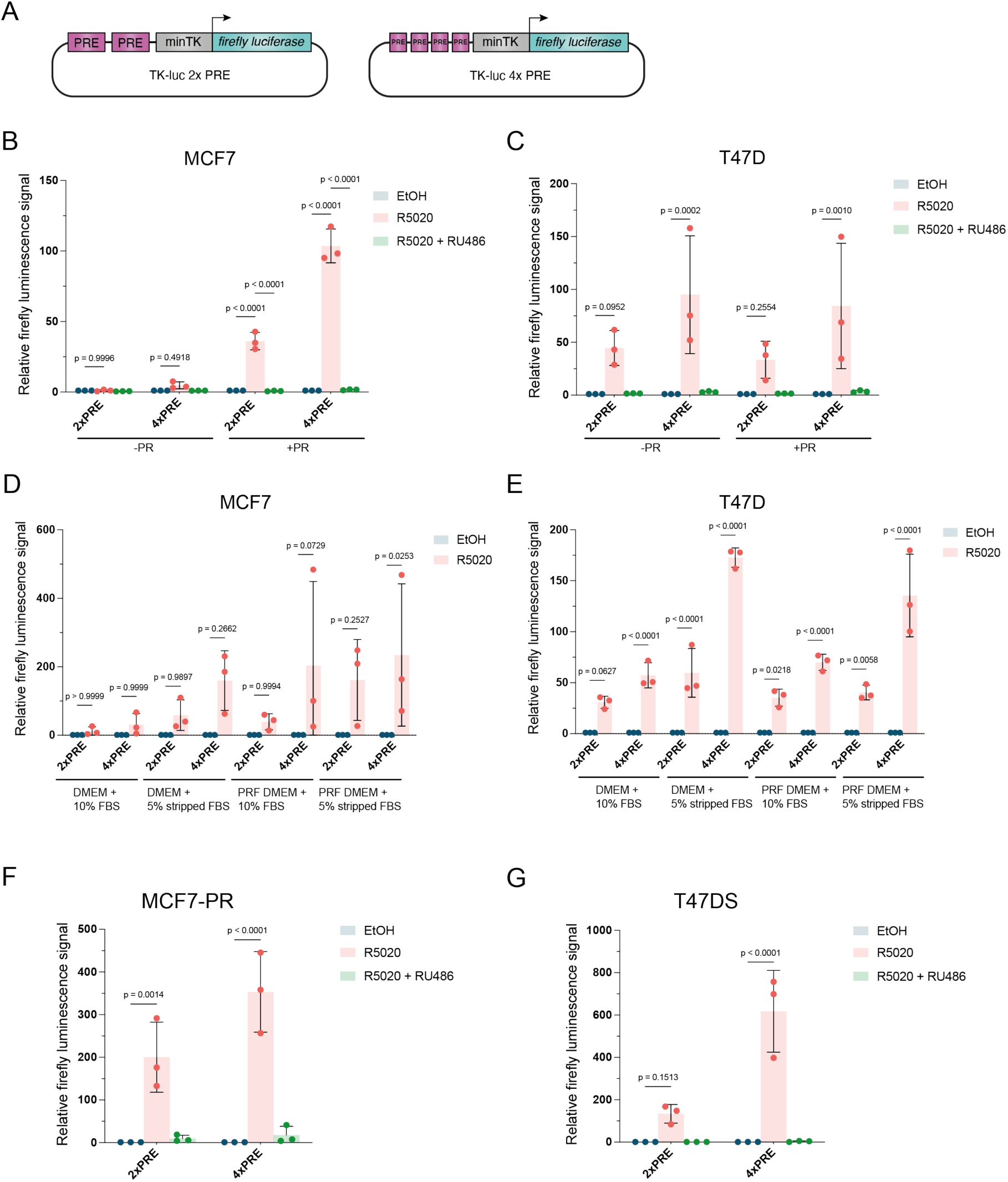
PR expression levels determine the strength of PRE-luciferase activity in MCF7 and T47D cells. a) Schematic representation of PRE constructs used in our dual luciferase assays. b-c) Relative luciferase activity of 2xPRE and 4xPRE-luciferase reporter constructs in MCF7 (b) or T47D (c) treated with EtOH (control, blue), 20nM R5020 (pink), or 20nM R5020 and 100nM RU486 (green) with (+) or without (-) PR co-transfection. Treated values are normalized for each condition over its own EtOH control. Every datapoint represents one biological experiment (n=3). Bars show the mean of the replicates. Error bars represent the standard deviation (SD) of the biological replicates. d-e) Relative luciferase activity of 2x/4xPRE-luciferase constructs in indicated medium for MCF7 (+PR) (d) or T47D (e) treated with EtOH (control, blue), 20nM R5020 (pink). PRF = phenol red free. f-g) Relative luciferase activity of 2x and 4xPRE-luciferase constructs treated with EtOH (control, blue), 20nM R5020 (pink), or 20nM R5020 and 100nM RU486 (green) for MCF7-PR (f) or T47DS (g) cells. Treated values are normalized for each condition over its own EtOH control. Every datapoint represents one biological experiment (n=3). Bars show the mean of the replicates. Error bars represent the standard deviation (SD) of the biological replicates.

For routine passaging, we cultured cells in phenol-red containing DMEM supplemented with 10% FBS. Since serum contains hormones [29], phenol red is known to be a weak ER agonist [30] and *PGR* is a known ER target gene [10], we reasoned that this might increase the baseline activity of our PRE reporters (and thus the background signal of our reporter assays) either directly or indirectly. To further improve the dynamic range of our PRE reporter assays, we therefore optimized the experimental medium conditions. In both MCF7 and T47D the use of charcoal stripped serum and phenol red free medium either alone or in combination, improved the signal to noise ratio of our 2xPRE and 4xPRE reporters (Fig2d,e). Because our MCF7 cells were less viable in phenol-red free medium, which is in line with the fact that they are known to be ER dependent [31], we chose regular phenol-red containing DMEM supplemented with 5% stripped FBS as the medium for all subsequent experiments.

We next aimed to obtain MCF7 and T47D cell lines that stably expressed high levels of PR. For MCF7, we integrated a lentiviral PR overexpression construct to create MCF7-PR. For T47D, we obtained a subclone that had previously been selected for high PR expression, named T47DS [32]. When both cell lines were assayed for *PGR* expression levels, the MCF7-PR cell line showed ∼6.5-fold higher *PGR* expression than our original MCF7 cell line, whereas T47DS expressed *PGR* at ∼3-fold higher levels than the parental T47D (SupFig1). As hypothesized, MCF7-PR and T47DS also showed a higher maximum luciferase signal compared to MCF7 and T47D (up to 350-fold and 600-fold induction of the 4xPRE reporter by R5020, respectively (Fig2f,g).

### Endogenous PR target gene inducibility correlates with PR expression levels

Having generated and selected MCF7 and T47D cell lines and culture conditions that are specifically suited for probing PR-mediated signaling questions, we next investigated if and how our readouts with exogenous reporter constructs translated to endogenous target gene induction. The literature reports variable and unreliable induction of presumed direct PR target genes in PR-expressing cell lines in response to progesterone or R5020 stimulation [15,17]. We therefore selected three known PR target genes (*WNT4* [33], *RANKL* [34], and *FKBP5* [18]) for qRT-PCR analysis. R5020 treatment of our parental MCF7 cells did not significantly induce expression of either endogenous *WNT4*, *RANKL*, or *FKBP5* (Fig3a, -PR). Transient overexpression of PR in MCF7 allowed us to measure induction of *RANKL* (∼2.2 fold) and *FKBP5* (∼3-fold), but not *WNT4* (Fig 2a, +PR). In T47D the induction of *RANKL* (4- vs 3-fold) and *FKBP5* (both ∼5-fold) was comparable regardless of whether PR was transiently overexpressed or not, while *WNT4* was not induced in either setting (Fig3b).

**Fig.3.**
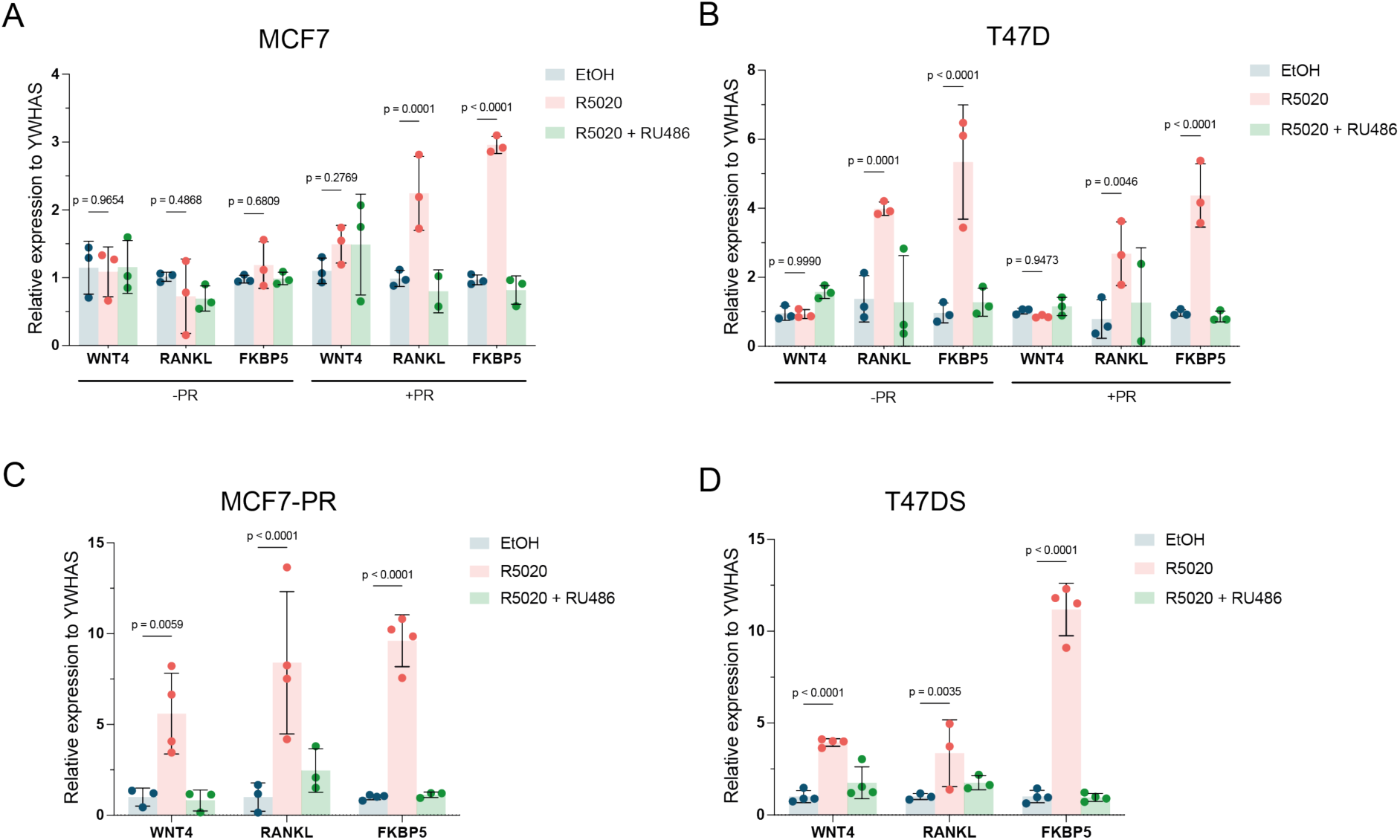
Induction of endogenous PR target genes in cell lines with different PR expression levels. a-b) *WNT4*, *RANKL* and *FKBP5* expression measured by qRT-PCR after EtOH (control, blue), 20nM R5020 (pink), or 20nM R5020 and 100nM RU486 (green) of MCF7 (a) or T47D (b) cells with (+) or without (-) PR co-transfection. Treated values are normalized for each condition over the mean EtOH control. Every datapoint represents one biological experiment (n=3). Bars show the mean of the replicates. Error bars represent the standard deviation (SD) of the biological replicates. c-d) *WNT4*, *RANKL* and *FKBP5* expression measured by qRT-PCR after EtOH (control, blue), 20nM R5020 (pink), or 20nM R5020 and 100nM RU486 (green) of MCF7-PR (c) or T47D (d) cells. Treated values are normalized for each condition over the mean EtOH control. Every datapoint represents the average from a technical duplicate of one biological experiment (n=4 total). Bars show the mean and error bars the standard deviation (SD) of the biological replicates.

In MCF7-PR cells, we measured more prominent induction compared to transiently transfected MCF7 cells, with *RANKL* (8-fold) and *FKBP5* (10-fold), as well as *WNT4* (5-fold) showing increased induction in response to R5020 treatment (Fig3c). In T47DS cells, which have the highest levels of *PGR* expression (SupFig1), *FKBP5* (∼11-fold) and WNT4 (∼4-fold) but not *RANKL* (∼3-fold) were induced to a better extent than in T47D (Fig3d). Taken together, an overall positive correlation exists between PR expression levels and the strength of the PR signaling response. PR levels are limiting in MCF7, for both the induction of PRE luciferase reporters (Fig2) and endogenous target genes (Fig3).

### PRE sequence variation to study PR signaling specificity and sensitivity

It is known that varying the absolute number of transcription factor binding sites affects the dynamic range of reporter gene constructs [34,35]. Therefore, we further modified our PRE-luciferase reporter by concatemerizing up to 12 PRE binding sites. Starting with the 4xPRE construct, we successively inserted two additional PRE sites via stepwise cloning, resulting in 6xPRE, 8xPRE, 10xPRE and 12xPRE reporter constructs. Contrary to our expectations, a clear peak in the luciferase response was observed for the 4xPRE luciferase reporter construct in both MCF7 and T47D (Fig4a,b). We therefore continued to use the 4xPRE reporter in subsequent experiments as our optimal PRE construct (Fig4a,b).

**Fig.4.**
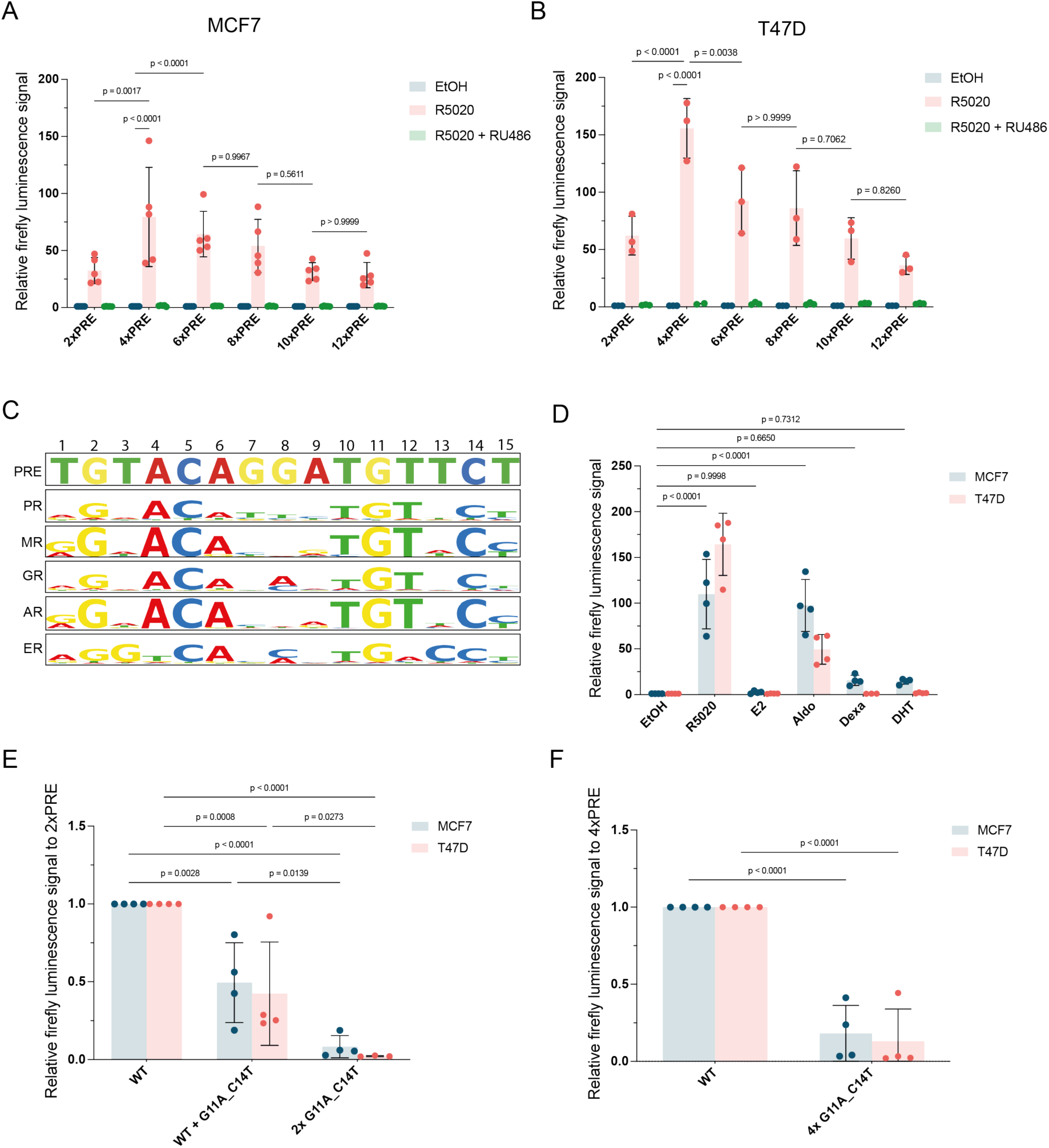
PRE sequence variations reveal PR signaling specificity and sensitivity. a-b) Relative firefly luciferase signal of 2-12xPRE-luciferase constructs after treatment with EtOH (control, blue), 20nM R5020 (pink), or 20nM R5020 and 100nM RU486 (green) of MCF7 (+PR) (a) or T47D (b) cells. Treated values are normalized for each condition over its own EtOH control. Every datapoint represents the average of technical duplicates from one biological experiment (n=3-5 total). Bars show the mean and error bars the standard deviation (SD) of the biological replicates. c) Visualization of the similarities between steroid nuclear receptor consensus sequences and the PRE consensus sequence in the luciferase constructs. d) Relative firefly luciferase signal of 4xPRE-luciferase construct after treatment of 10nM R5020, 10nM E2, 10nM aldosterone, 10nM dexamethasone or 10nM dihydroxytestosterone in MCF7 (+PR) (blue) or T47D (red) cells. Treated values are normalized for each condition over its own EtOH control. Every datapoint represents the average of technical duplicates from one biological experiment (n=4 total). Bars show the mean and error bars the standard deviation (SD) of the biological replicates. e-f) Relative firefly luciferase signal of 2xPRE (e) or 4xPRE-luciferase (f) constructs with G11A_C14T variation in one of the two or both PRE sites (2xPRE) (e) or all four sites (4xPRE) (f) after 20nM R5020 treatment of R5020 in MCF7 (+PR) (blue) or T47D (red) cells. Treated values are normalized for each condition over the WT induction. Every datapoint represents the average of technical duplicates from one biological experiment (n=4 total). Bars show the mean and error bars the standard deviation (SD) of the biological replicates.

The PRE site in our reporter is derived from the consensus PRE [21,36]. While ER binds to specific ER response elements (ERE), PR shares its consensus sequence with other steroid hormone receptors, including the mineralocorticoid receptor (MR), glucocorticoid receptor (GR) and androgen receptor (AR) (Fig4c) [38]. Both T47D and MCF7 have been reported to express at least some of these additional nuclear hormone receptors [19,37]. Moreover, PR and GR were previously shown to activate transcription by interacting with the same responsive element [38,39].

To test the response of our 4xPRE luciferase construct to other steroid hormone signals, we performed luciferase assays in MCF7 and T47D following stimulation of the cells with β-estradiol (E2, the ER ligand), aldosterone (Aldo, the MR ligand), dexamethasone (Dex, the GR ligand) or dihydroxytestosterone (DHT, the AR ligand) in comparison to R5020. As before, R5020 induced PRE-reporter signal ∼100-fold in MCF7 (+PR) and ∼150-fold in T47D. Out of all tested non-progesterone ligands, only aldosterone was able to significantly activate the reporter in MCF7 (∼100-fold) as well as in T47D (∼50-fold) (Fig4d). A minor increase was observed for dexamethasone and dihydroxytestosterone. Thus, our 4xPRE-reporter can also be activated by related receptors, but only in a setting where they are specifically activated instead of PR.

Whereas our luciferase constructs contain consensus PR binding sites, *in vivo* enhancers have been reported to use suboptimal transcription factor binding sites to ensure specific and robust gene regulation [41,42]. For example, imperfect estrogen responsive elements (EREs) are important for ER binding and synergize with perfect EREs [44]. In addition, endogenous progesterone-responsive enhancers have been described to contain imperfect PREs [38]. Therefore, an as of yet unanswered question in the PR field is what the effect is of natural variations in DNA sequences on PR binding and, subsequently, on progesterone-dependent gene regulation.

We reasoned that we should now be able to quantitatively measure the biological activity of PRE sequence variations at the single nucleotide level to start solving this question. We took two variations of the PRE consensus sequence (G11A and C14T) that had both previously been described to abolish PR binding to the PRE sequence as well as to impair biological PRE activity [37]. We generated 2xPRE-luciferase constructs containing one consensus PRE and one PRE with the G11A and C14T base pair variations (Fig4e). A double G11A and C14T mutation in a single PRE site causes a statistically significant, ∼2-fold reduction in luciferase signal in MCF7 upon R5020 stimulation (Fig4e). We hypothesized that remaining activity likely represented activity of the remaining wildtype PRE. Introducing the G11A and C14T mutations into both sites of the 2xPRE reporter or all four sites of the 4xPRE luciferase construct, indeed resulted in a ∼92% or ∼82% (MCF7) and a ∼98% or ∼87% (T47D) loss of progesterone-inducibility, respectively (Fig4e,f). Taken together, mutations in the consensus sequence modulate activity of the PRE site and can therefore be used to probe the effects of sequence variations.

### PRE-GFP reporter constructs to visualize of PR signaling in individual cells

As stated, nuclear PR protein localization does not necessarily reflect signaling status [6,22, 23] (Fig1). To be able to resolve PR signaling activity at the single cell rather than the population level, we generated lentiviral PRE-GFP constructs with 2, 4 or 6 consensus PRE sites. We stably introduced these constructs into MCF7-PR cells and FACS sorted a polyclonal population that induced GFP in response to R5020 treatment (Fig5a).

**Fig.5.**
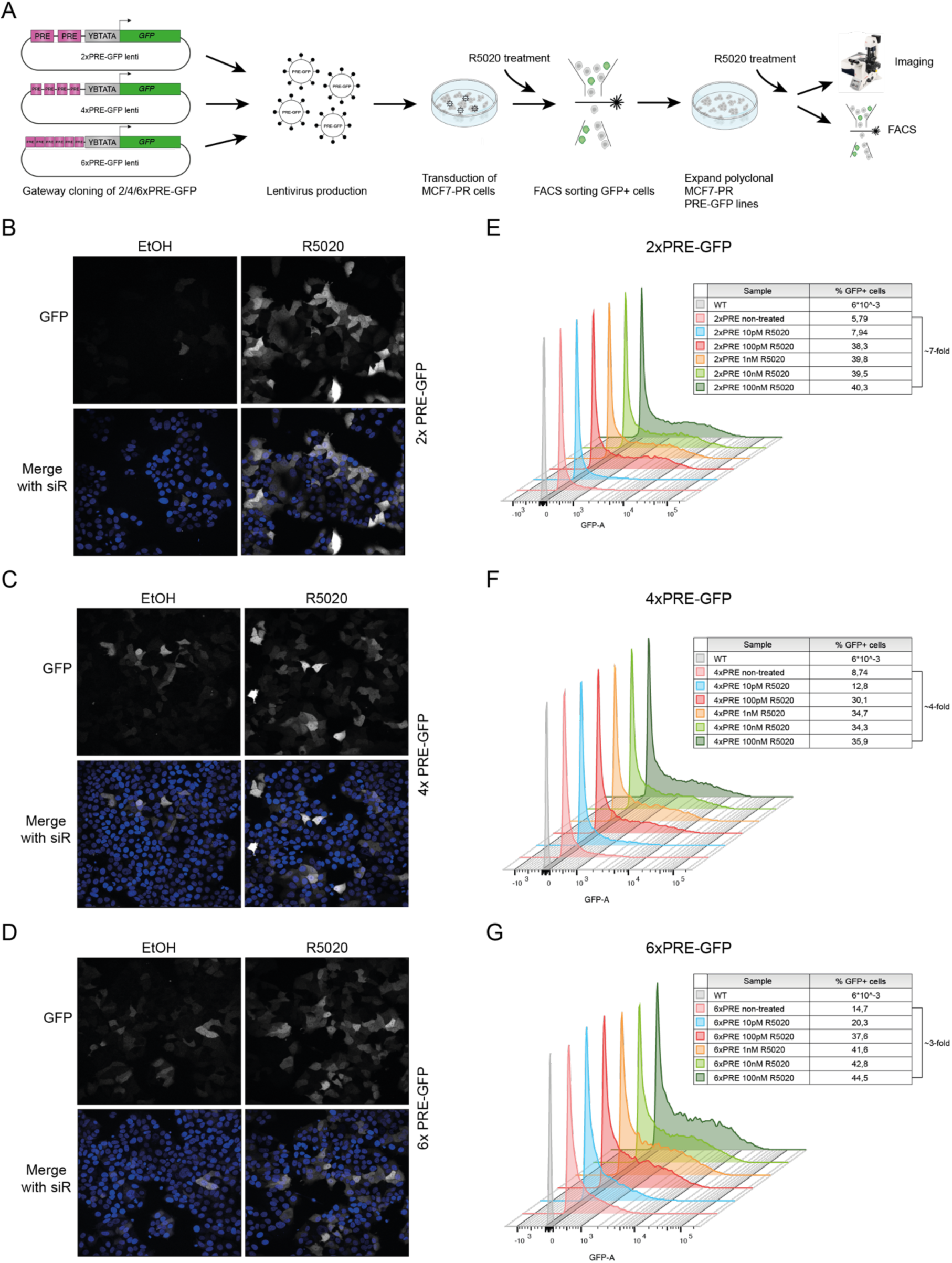
PRE-GFP reporter cell lines as a tool to visualize PR signaling at the single cell level. a) Schematic representation of the experimental workflow for generating 2x, 4x, and 6xPRE-GFP MCF7-PR lines. b-d) Representative confocal microscope images of 2x (b), 4x (d) or 6xPRE-GFP (d) MCF7-PR cells treated with EtOH as a control or treated with 20nM R5020 for 22 hours from n-=2 experiments. Nuclei were counterstained with SiRDNA (blue). e-g) Representative histograms depicting FACS analysis of 2x (e), 4x (f) or 6xPRE-GFP (g) MCF7-PR cells treated with indicated treatments for 22 hours of n=2 experiments. Tables show the percentage of GFP+ cells for each treatment.

The resulting 2x/4x/6xPRE-GFP cell lines were further characterized by imaging and FACS. R5020 treatment induced the GFP reporter over background in all three cell lines as visualized by fluorescence microscopy (Fig5b-d). However, the response was heterogeneous, as not all cells responded despite the fact that they had been FACS sorted for their ability to do so. GFP intensity also varied between individual cells. To determine sensitivity of the reporters, we treated the 2x/4x/6xPRE-GFP lines with an R5020 concentration range from 10pM-100nM and quantified the percentage of GFP positive cells by FACS analysis. We observed a modest induction of all reporter constructs even in response to 10pM R5020, which is in the physiological range (Fig5e-g). The maximum response was reached in response to 100pM R5020 (Fig5e-g, ∼38% for 2xPRE vs. 5.8% GFP+ in the untreated cells, ∼30% for 4xPRE vs. 8.7% GFP+ in the untreated cells, ∼38% for 6xPRE vs. 14.7% GFP+ in the untreated cells). R5020 concentrations exceeding 100pM did not substantially increase the percentage of GFP positive cells any further in either of the cell lines. In conclusion, our lentiviral 2x/4x/6xPRE-GFP reporters display robust GFP inducibility and are capable of detecting PR activity at physiological concentrations, which should also make them suitable for *in vivo* applications.

### R5020 dose-response measurements reveal PR signaling dynamics

Progesterone concentrations in blood range from the low picomolar range during menopause to the micromolar range during pregnancy [2,9]. For experimental *in vitro* studies, cells are routinely treated with 10-100nM R5020 [45]. We therefore used 20nM R5020 for most of our experiments depicted in Figure 1-4. However, since R5020 is a synthetic progestin that is more stable and has a higher intracellular availability than progesterone, these concentrations are at the higher end of the physiological progesterone range. To our knowledge, how PR signaling responds to different doses of ligand has not been studied extensively. Having already been able to measure induction of our PRE-GFP reporter in response to treatment with 10pM R5020 (Fig5e-g), we therefore also examined the response of our PRE luciferase reporter constructs as well as our selection of endogenous PR target genes to a range of concentrations of R5020 in both MCF7 and T47D cells.

Interestingly, stimulation with 1pM to 100nM R5020 revealed distinct response patterns. First, dose-response curves were quite comparable for the 2x, 4x and 6x PRE-luciferase reporter constructs (SupFig2a,b) but differed between MCF7 and T47D (Fig6a). Specifically, in MCF7 the PR signaling response peaked at a much lower concentration (10pM R5020) than in T47D (1nM R5020). Second, R5020 concentrations higher than 1nM do not significantly increase the PR signaling response in either cell line, suggesting that saturation is reached at R5020 levels that are lower than typically used in the literature (Fig6a). Third, endogenous target gene expression varied substantially depending on both the target and the cell line (Fig6b-d, SupFig2c,d). As before, *WNT4* expression was significantly induced in MCF7 but not in T47D (Fig 3a,b, Fig6b), with *WNT4* levels increasing in response to stimulation with 100pM R5020 and with higher concentrations not improving the response. *RANKL* expression did not show robust induction under any condition (Fig6c). Finally, the negative feedback target gene *FKBP5* was robustly and dose-dependently expressed in both MCF7 and T47D starting at a concentration of 100pM R5020 (Fig6d). Together, these results show the importance of matching the concentration of the hormone stimulus to the cell line and the experimental readout.

**Fig.6.**
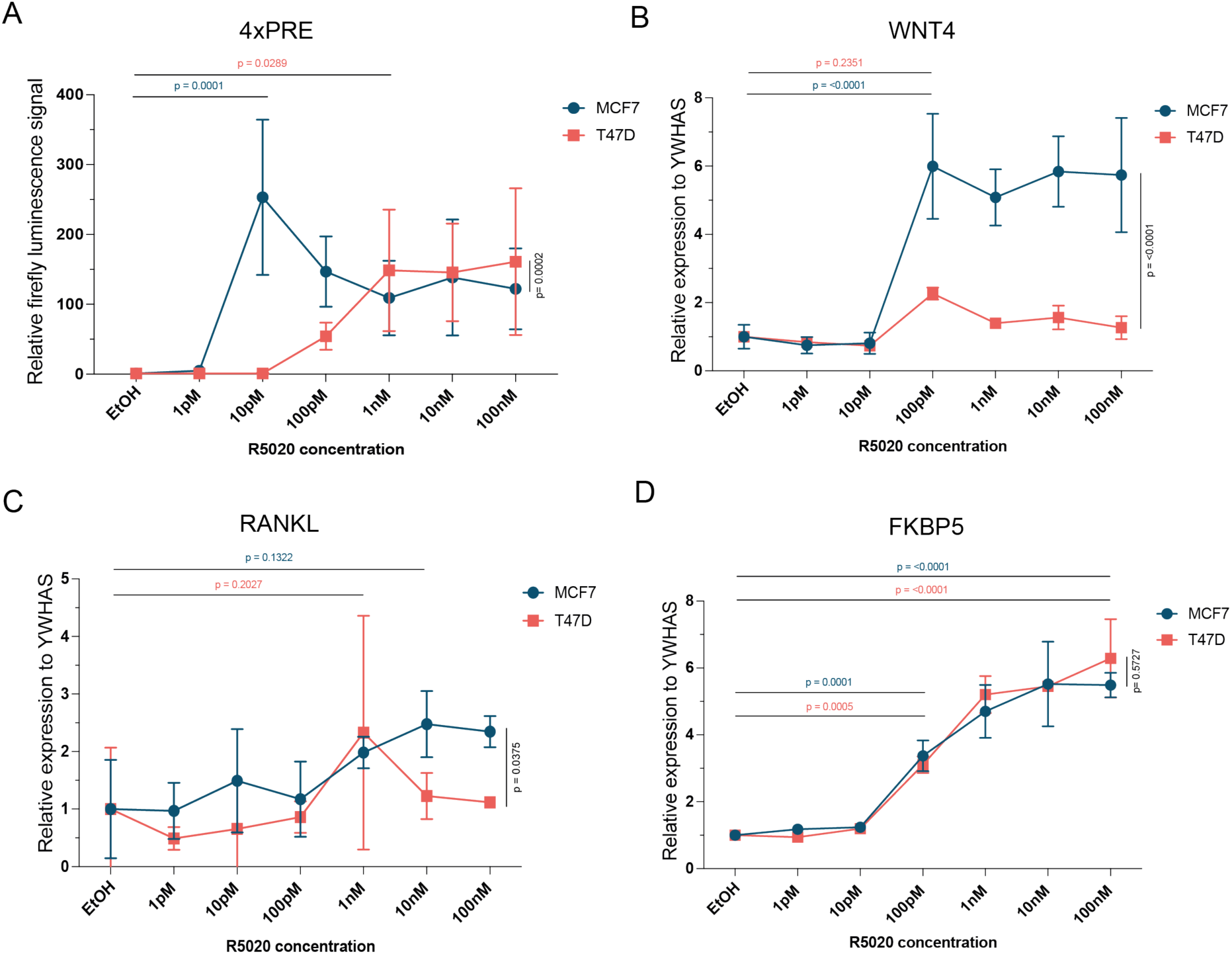
PR signaling response varies per cell line, target gene and R5020 concentration used. a) Relative firefly luciferase signal of 4xPRE-luciferase after indicated R5020 treatments in MCF7 (+PR) (blue) or T47D (red) cells. Treated values are normalized for each condition over the mean EtOH control. Every point represents the mean of n=3 biological experiments. Error bars represent the standard deviation (SD) of the biological replicates. b-d) *WNT4* (b), *RANKL* (c) and *FKBP5* (d) expression measured by qRT-PCR after indicated R5020 treatments in MCF7 (+PR) (blue) or T47D (red) cells. Treated values are normalized for each condition over the mean EtOH control. Every point represents the mean of n=3 biological experiments. Error bars represent the standard deviation (SD) of the biological replicates.

## Discussion

PR signaling is of fundamental importance for breast development and physiology, but it remains understudied in both the healthy breast and in breast cancer. One bottleneck has been the availability of reliable readouts to measure PR signaling responses in breast epithelial cells. Here, we describe a toolbox for quantitative analyses of PR signaling, which we test in the widely used MCF7 and T47D breast cancer cell lines. We show that the absolute PR protein levels determine the strength of the PR signaling response (Fig2). This could shed new light on previous studies that failed to detect the expected patterns of PR target gene induction [7]. After optimizing the culture media (Fig2) and R5020 treatment (Fig6) conditions, we find that both PRE luciferase reporter assays and qRT-PCR analysis of endogenous PR target genes can be useful readouts for quantifying PR signaling activity (Fig2, Fig3, Fig6).

Our new 4xPRE luciferase reporter construct has a high dynamic range, and we therefore recommend it for robust and reliable measurements of PR signaling. Care should be taken under conditions where related nuclear hormone receptor signaling pathways may be active, since our PRE consensus site can also be activated by MR in response to aldosterone stimulation (Fig4). We expect this reporter to serve as an improved starting point, compared to previous analysis [37], for studying the impact of PRE site variations, for example in the context of suboptimal PRE sites that are likely to be present in PR-dependent enhancers *in vivo*. In addition to varying the sequence, changing the order, total amount, orientation and spacing of PRE sites may also provide meaningful new insights into PR-PRE binding and PR signaling dynamics.

We show that nuclear accumulation of PR is independent of ligand-activated PR (Fig1). Thus, nuclear localization does not equal PR signaling activity. We present our lentiviral PRE-GFP reporter constructs as an alternative, functional readout and a new tool to visualize PR signaling activity at the single cell level. Applying PRE-GFP on top of PR staining in future experiments can confirm PR signaling activity and thus provide a key additional readout in cultures containing PR positive cells. It should be noted that we observe a heterogeneous response in our tested PRE-GFP cell lines, with a maximum of 40% of cells responding. This could be due to technical reasons, since we analyzed polyclonal cell populations in which the lentiviral PRE-GFP cassette may have integrated in different genomic locations and chromatin contexts that may be more or less conducive to expression and PR-mediated induction. Alternatively, this result could reflect the underlying biology as it is still unknown if and how PR-A and PR-B expression levels or PR signaling can fluctuate over time and across cell populations. As it is known that PR requires phosphorylation on several residues for its proper activation and downstream effects, the interplay or abundance between PR and its activating kinases (such as MAPK and CDK2) could be one of the explanations for the heterogeneous response [45,46]. Also, cell cycle stage could affect the PR signaling outcome [48]. Thus, at least some cell-to-cell heterogeneity in the response is to be expected (Fig 1).

One outstanding question in the field is how PR signaling, induced by experimental – and typically high – doses of R5020 compares to more physiological doses of progesterone. Recently, it was reported that breast cancer cells are able to react to physiological progestin concentrations (50pM) [45]. We confirmed the response of our PRE-GFP and PRE-luciferase constructs as well as two endogenous PR target genes (*WNT4* and *FKBP5*) to physiologically relevant concentrations of R5020 (Fig5, Fig6), although the response does differ per cell line and target gene. For example, *RANKL* expression did not consistently show statistically significant induction (Fig2, Fig6c), although increased *RANKL* expression was observed in some individual experiments. We suspect that this is due to low absolute levels of baseline *RANKL* expression levels in MCF7 and T47D cells, resulting in high standard deviations and variable induction. Thus, in our hands *WNT4* and *FKBP5* are more robust PR targets compared to *RANKL*.

One of the conceivable explanations for our measured differences in PR target gene activation across different cell lines could be that PR expression levels play a role here as well. PR expression is important for PR binding site accessibility, as lowering PR expression in T47D cells exhibited a decrease in accessibility of PR binding sites [45]. This phenomenon could explain our observed differences in target gene expression in MCF7 WT, PR transfected and stable PR cells and T47D vs. T47DS cells (Fig2). Furthermore, it is a possibility that PR-A/PR-B ratios play a role (SupFig1), since PR-A/PR-B ratios effect the transcriptional outcome of PR signaling [11,48].

Summarizing, we present new approaches for measuring PR signaling in breast epithelial cells. In our hands, a 4xPRE luciferase reporter is the most sensitive and reliable tool to measure PR signaling activity. Endogenous target gene expression analysis is still a valuable addition, with the negative feedback target *FKBP5* serving as the most robust PR target gene. However, it is recommended to take multiple target genes along as their response and level of induction will differ depending on the cellular context. Additionally, our PRE-GFP constructs are a valuable tool for single cell PR signaling visualization and measurements. We expect these new tools and optimized conditions to be a useful foundation for addressing outstanding questions regarding the molecular mechanisms that determine the strength and dynamics of PR signaling in mammary gland biology.

**Supplemental Fig.1.**
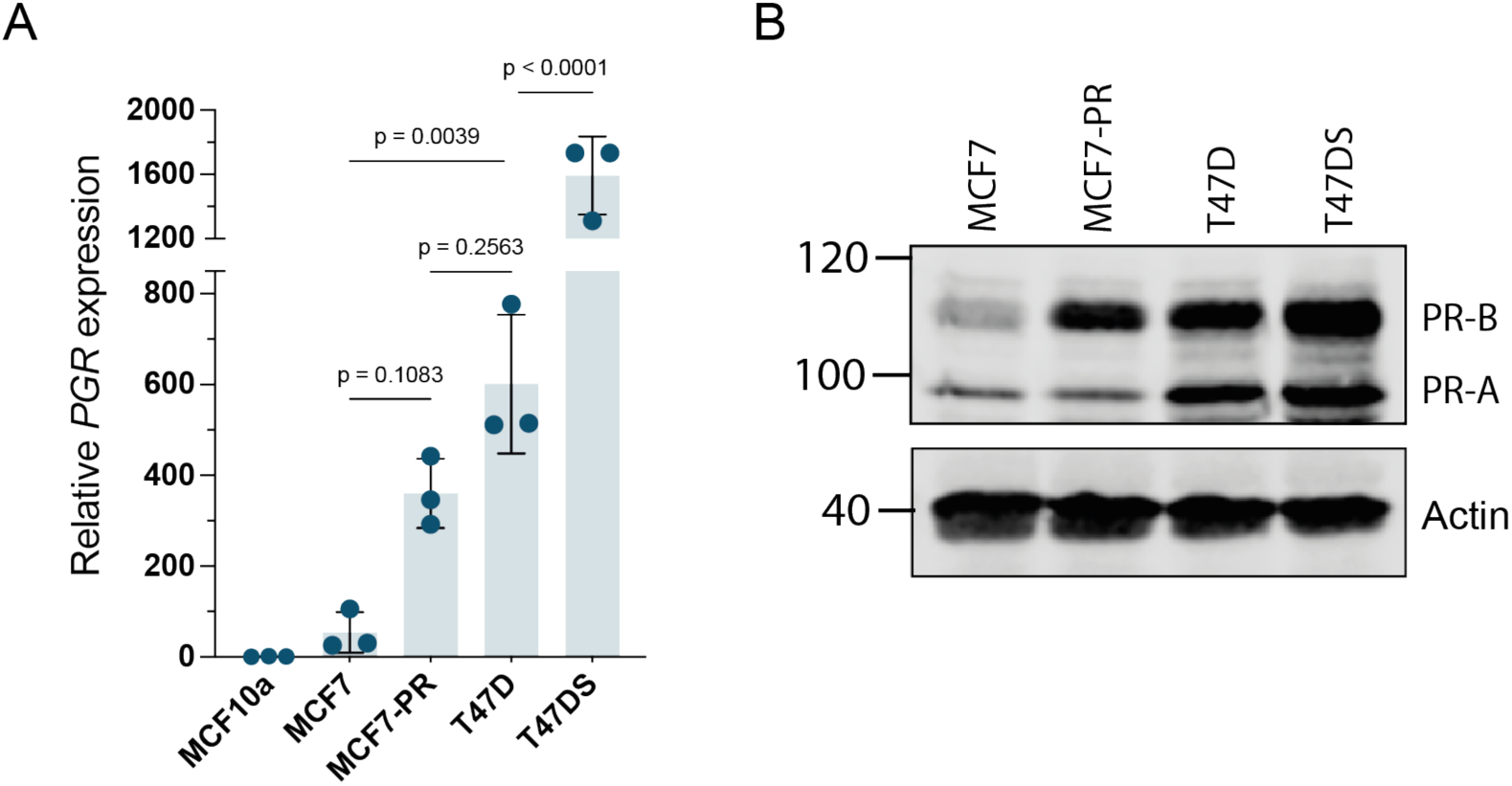
PR expression levels of MCF7, MCF7-PR, T47D and T47DS cell lines. a) *PGR* expression of MCF7, MCF7-PR, T47D and T47DS cells as determined by qRT-PCR. YWHAS was used as a reference gene, *PGR* expression values were normalized to MCF10a (n=3 biological replicates). Data points indicate the individual biological replicates, and the bar graphs the mean relative *PGR* expression b) Western blot showing PR expression in MCF7, MCF7-PR, T47D and T47DS cells. Although a full-length PR construct is overexpressed in MCF7-PR, this results mainly in expression of the PR-B isoform.

**Supplemental Fig.2.**
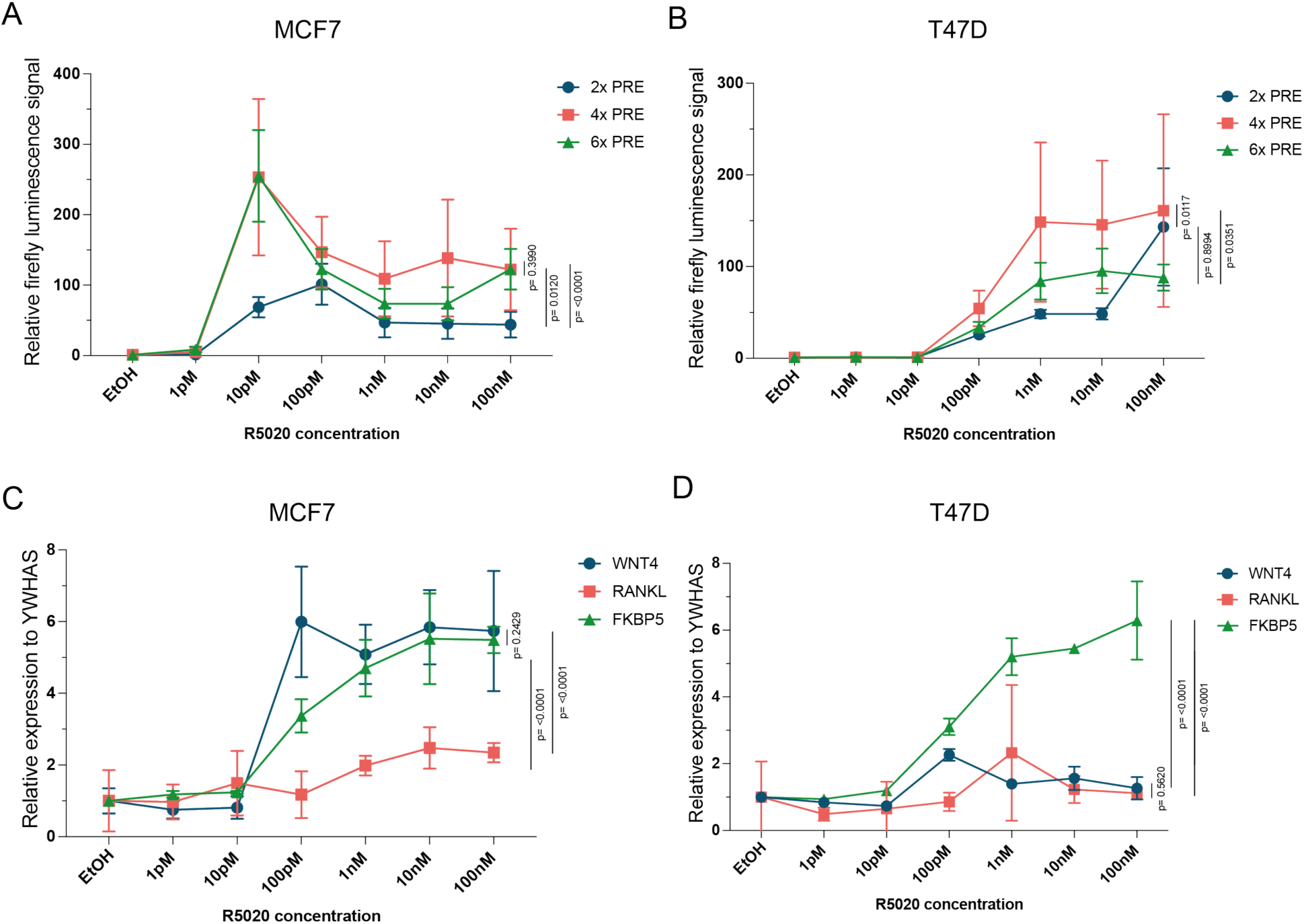
Response differences to increasing R5020 concentrations between luciferase constructs, target genes and cell lines. a-b) Relative firefly luciferase signal of 2x, 4x and 6xPRE-luciferase after indicated R5020 treatments MCF7 (+PR) (a) or T47D (b) cells. Treated values are normalized over the mean EtOH control. 4xPRE results are the same as depicted in Fig6a. c-d) *WNT4*, *RANKL* and *FKBP5* expression measured by qRT-PCR after indicated R5020 treatments. Treated values are normalized for each condition over the mean EtOH control. Every point represents the mean of three biological experiments. Error bars represent the standard deviation (SD) of the biological replicates. Results are the same as depicted in Fig6b-d but are here sorted on cell line instead of target gene.

## Material and methods

### Cell culture

Human MCF7 breast cancer cells (a kind gift from Prof. Dr. Pernette Verschure, Swammerdam Institute for Life Sciences, Amsterdam, The Netherlands), Human T47D and T47DS breast cancer cells (a kind gift from Dr. Stieneke van den Brink, Hubrecht institute, Utrecht, The Netherlands), and HEK293TN cells (System Biosciences, #LV900A-1) were routinely cultured in Dulbecco’s Modified Eagle Medium (DMEM) containing GlutaMAX (Gibco, #11584516), supplemented with 10% Fetal Bovine Serum (FBS) (Thermo Fisher, #11573397). Cells were split 1:10 every 3-4 days and routinely tested for mycoplasma. MCF7 cells transfected with a PR expression construct are referred to as MCF7 (+PR). MCF7 cells stably expressing a lentiviral PR construct are referred to as MCF7-PR.

### Immunofluorescence staining and fluorescence microscopy

One day prior to imaging, 75,000 MCF7 or 100,000 T47D cells were seeded on an 8 well chamber slide with glass bottom (Ibidi, #80827-90). The next day, cells were treated with 20nM R5020 (Promegestone, Perkin Elmer, #NLP004005MG), or 20nM R5020 and 100nM RU486 (Mifestrone, Sigma Aldrich, #475838) dissolved in ethanol, which was also taken along as a negative control. Two hours after treatment, the cells were fixed in 4% paraformaldehyde (Alfa Aesar, #43368) in PBS for 15 minutes at room temperature (RT) and washed with HBSS (Thermo Fisher, #11550456) three times. Next, the cells were permeabilized with 0.1% Triton in PBS for 15 minutes at RT. After three washes with HBSS, the samples were blocked for 2 hours in HBSS with 4% BSA (Tocris BioScience, #5217). Incubation with primary antibody rabbit anti progesterone receptor (1:500 in 4% BSA, Cell Signaling #8757/D8Q2J, recognizing both PR-A and PR-B) was performed overnight (O/N) at 4°C. Following three washes with HBSS, the cells were incubated for 2 hours with secondary antibody AlexaFluor 488 Goat Anti-Rabbit IgG (1:1000 in 4% BSA, Invitrogen #A11008) at RT in the dark. The samples were stained with DAPI (1:1000 in HBSS, Invitrogen, #D1306) for 10 minutes at RT, washed three times with HBSS and imaged on an SP8 confocal microscope (Leica Microsystems). Imaging was performed using a 63x oil objective with 405 (5% laser power) and 488 (8% laser power for MCF7 and 2% laser power for T47D) lasers, using a PMT1 detector (gain 700) for fluorescent signal with a 413-469 bandpass for DAPI and a HyD detector (gain 100) for fluorescent signal with a 496-547 bandpass for GFP. Images for the individual channels were extracted using Fiji [50]. Care was taken to image each sample with the same settings. For image analysis, Cell Profiler [51] was used to segment the nucleus based on the DAPI signal, excluding border nuclei. The nuclear mean intensity of the PR staining was then measured for each individual cell, normalized to the average intensity in the non-treated condition and plotted in GraphPad Prism (Version 10.0.0).

### DNA cloning

The 2xPRE-luciferase reporter (2X PRE TK luc) construct contains 2 consensus PRE sites upstream of a minimal thymidine kinase (TK) promoter (Table 2, #1). The pcDNA3-PRB plasmid contains the full-length PR sequence (Table 2, #2). To generate PRE-luc constructs containing concatemerized, wildtype or mutated PRE sites, primers were designed using Snapgene (Table 1). The PRE-luc constructs were generated using restriction cloning of the annealed oligonucleotides into the 2xPRE luc vector with BamHI (Thermo fisher, #FD0054). For the vectors with multiple PRE sites, restriction enzyme digestion and ligation were repeated until constructs with 4, 6, 8, 10 and 12x PRE sites were obtained.

**Table 1:**
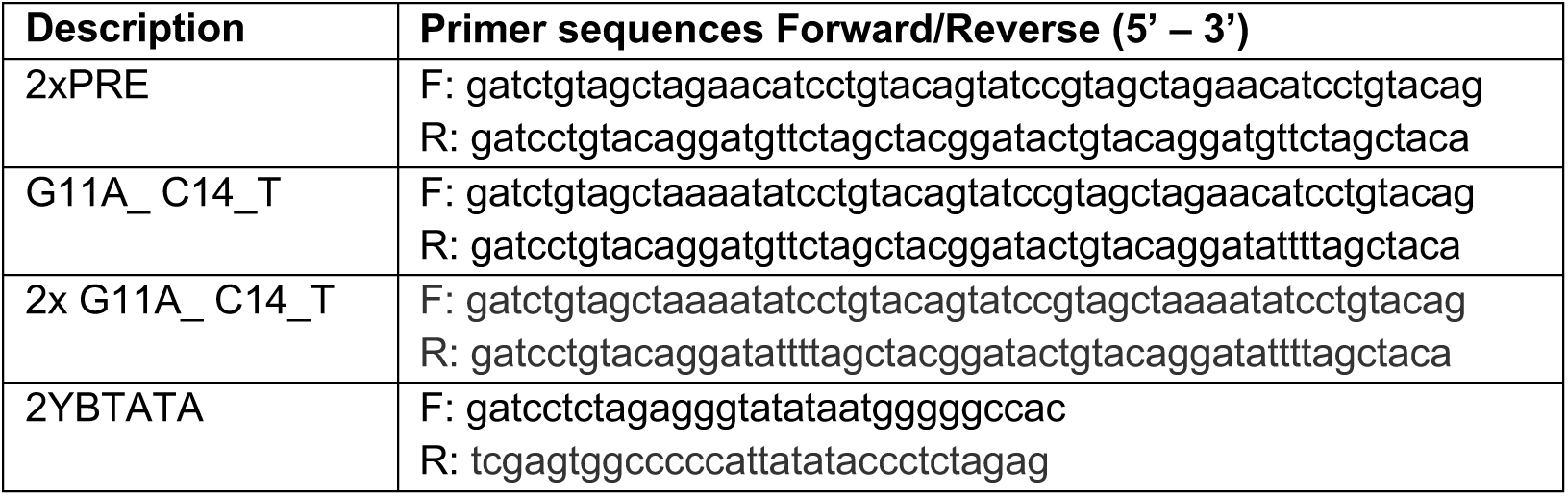
Primer sequences.

**Table 2:** Used Plasmids.

| Plasmid # | Plasmid name | A gift from | Ref | Addgene number |
| --- | --- | --- | --- | --- |
| 1 | 2X PRE TK luc | Donald McDonnell | [22] | #11350 |
| 2 | pcDNA3-PRB | Elizabeth Wilson | [54] | #89130 |
| 3 | pMuLE-ENTR-MCS-R4-R3 | Ian Frew | [55] | #62086 |
| 4 | pMuLE ENTR MCS L1-L4 | Ian Frew | [55] | #62087 |
| 5 | pMVP (L1-L4) CMV promoter | Christopher Newgard | [56] | #121686 |
| 6 | pMVP (L3-L2) polyA | Christopher Newgard | [56] | #121746 |
| 7 | pMVP/Lenti/Blast-DEST | Christopher Newgard | [56] | #121849 |
| 8 | pMVP (R4-R3) eGFP | Christopher Newgard | [56] | #121730 |
| 9 | pMVP/Lenti/Neo-DEST | Christopher Newgard | [56] | #121850 |
| 10 | pCMVDR8.2 | Bob Weinberg | [57] | #8455 |
| 11 | RSV-rev | Didier Trono | [58] | #12253 |
| 12 | VSVg | Didier Trono | - | #12259 |

For generation of lentiviral PR expression vectors, full-length PR was cloned into the multisite gateway compatible pMuLE-ENTR-MCS-R4-R3 (Table 2, #3) by restriction cloning from pcDNA3-PRB (Table 2, #2). For generation of entry clones containing 2x, 4x, or 6xPRE sites for multisite gateway reactions, annealed oligonucleotides coding for the 2YBTATA minimal promotor [52] (Table 1) were ligated into the pMuLE ENTR MCS L1-L4 vector (Table 2, #4), followed by ligation of annealed 2xPRE oligonucleotides (Table 1).

Lentiviral 2/4/6xPRE-GFP and CMV-PR plasmids were generated using multisite LR gateway reactions. Gateway vectors were diluted to 10 fmol/ul of each entry plasmid and 20 fmol/ul of the destination plasmid with TE buffer (pH 8.0). Lenti-CMV-PR constructs were created by mixing 10 fmol 5’ entry CMV promoter (Table 2, #5), 10 fmol pMuLE (R4-R3) PRB, 10 fmol 3’ entry polyA (Table 2, #6), and 20 fmol destination vector Lenti-Blast (Table 2, #7). For Lenti-2/4/6xPRE-GFP reporters: 10 fmol 5’ entry plasmids containing 2/4/6xPRE and 2YBTATA, 10 fmol middle entry eGFP (Table 2, #8), 10 fmol 3’ entry polyA (Table 2, #6), and 20 fmol destination vector Lenti DEST (Table 2, #9) were combined. LR Clonase II Plus enzyme (Thermo Fisher Scientific, #12538120) was added to catalyze the gateway reactions according to the manufacturer’s instructions.

### Dual luciferase assays

MCF7, MCF7-PR, T47D or T47DS cells were plated in 24-well plates at a density of 100,000 (MCF7/MCF7-PR) or 150,000 (T47D/T47DS) cells per well. After 24 hours, the medium was replaced with DMEM supplemented with 5% Charcoal Stripped Fetal Bovine Serum (Thermo fisher, #A3382101), or one of the following four media for optimization of medium conditions: 1) DMEM supplemented with 10% FBS (identical to the standard propagation media), 2) DMEM supplemented 5% Charcoal Stripped Fetal Bovine Serum (stripped FBS – the final media used for all experiments after Fig2C), 3) phenol red free (PRF) DMEM (Thermo Fisher, #11594416) supplemented with 10% FBS or 4) phenol red free DMEM supplemented with 5% Charcoal Stripped Fetal Bovine Serum. Cells were transfected using X-tremeGENE HP DNA Transfection Reagent (Sigma-Aldrich, #06366546001), according to the manufacturer’s instructions. For MCF7, transfections were performed in duplicate, using a total amount of 500ng plasmid DNA per well, consisting of 200ng Luciferase construct and 200ng PR expression vector (Table 2, #2) and 100ng Renilla construct. For MCF7-PR, T47D and T47DS 400ng Luciferase construct and 100ng Renilla construct was used. On the next day, cells were treated with either EtOH (control), 20nM R5020 (Perkin Elmer, #NLP004005MG), or 20nM R5020 and 100nM RU486 (Sigma Aldrich, #475838) in fresh medium. When comparing different R5020 concentrations, the cells were treated with the indicated range of R5020 (1pM – 100nM) or EtOH as the solvent control. To determine reporter specificity, the cells were treated with 10nM of the following compounds: R5020, β-estradiol (Sigma, #E8875-1G), aldosterone (Sigma-Aldrich, #A9477-5MG) dexamethasone, (Sigma-Aldrich, #D4902), dihydroxytestosterone (Merck, #D-073) or EtOH as solvent control. Cells were lysed after exactly 24 hours of stimulation in 1X Passive Lysis Buffer (100 μl per well (Promega, #E1941)). For the dual luciferase measurements, non-commercial firefly, and Renilla Luciferase Reagents (LAR) were used [53]. Firefly and Renilla luciferase activity was measured in a GloMax Navigator (Promega, #GM2000). Firefly luciferase values were normalized to Renilla luciferase values. The data is presented as fold-change in firefly luciferase activity, normalized over the non-treated control, unless stated otherwise. Plots are generated using GraphPad Prism. (Version 10.0.0).

### qRT-PCR

MCF7, MCF7-PR, T47D and T47DS cells were plated in 6-well plates at a density of 300,000 (MCF7/MCF7-PR) or 400,000 (T47D/T47DS) cells per well. The following day, for medium was refreshed for all cells and MCF7 cells were transfected with 2µg pcDNA3-PRB per well using X-tremeGENE HP DNA Transfection Reagent (Sigma Aldrich, #06366546001) according to the manufacturer’s instructions. 48 hours after plating, cells were treated with EtOH, 20nM R5020 (Perkin Elmer, #NLP004005MG), or 20nM R5020 and 100nM RU486 (Sigma Aldrich, #475838). When comparing different R5020 concentrations, the cells were treated with 1pM-100nM R5020 or EtOH. After 24 hours treatment, RNA was isolated using Trizol according to the manufacturer’s instructions. Total RNA was DNAse treated with RQ1 DNAse (Promega, #M6101). cDNA synthesis was performed using 4000ng RNA using SuperScript IV Reverse Transcriptase (Invitrogen, #18090200) and Random Hexamers (Invitrogen, #N8080127) according to manufacturer’s guidelines with the addition of RiboLock RNase Inhibitor (Thermo Scientific, #EO0328). cDNA was diluted 10-fold and qRT-PCR reactions were performed using 5x HOT FIREpol EvaGreen qPCR mix plus (ROX) (Bioconnect, #08-24-00020) and a QuantStudio 3 Real-Time PCR System (Thermo Fisher). qRT-PCR primers were thoroughly checked to have a single melt curve. Primer sequences are listed in Table 3. Calculations were performed using the ddCt method and presented as relative values normalized over YWHAS and EtOH treated conditions. Plots are generated using GraphPad Prism (10.0.0).

**Table 3:**
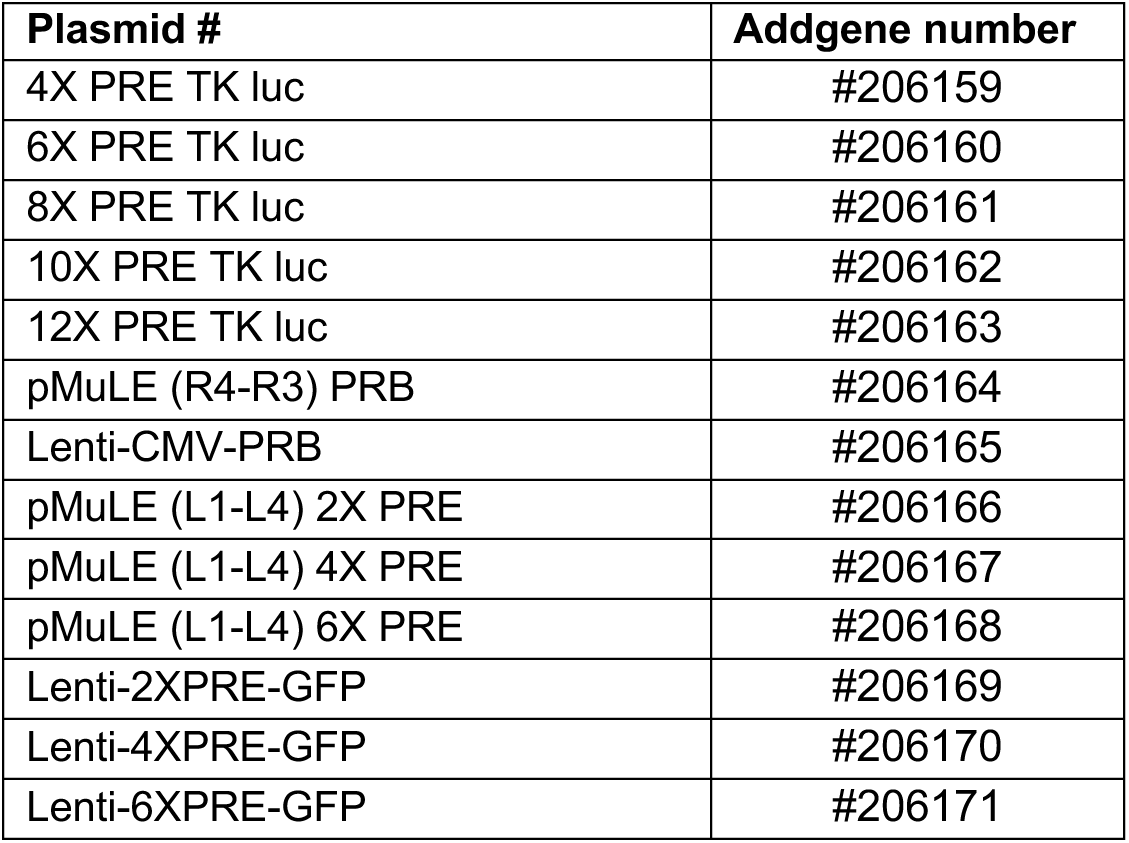
Generated plasmids. (Addgene deposition in progress at the time of submission)

**Table 4:**
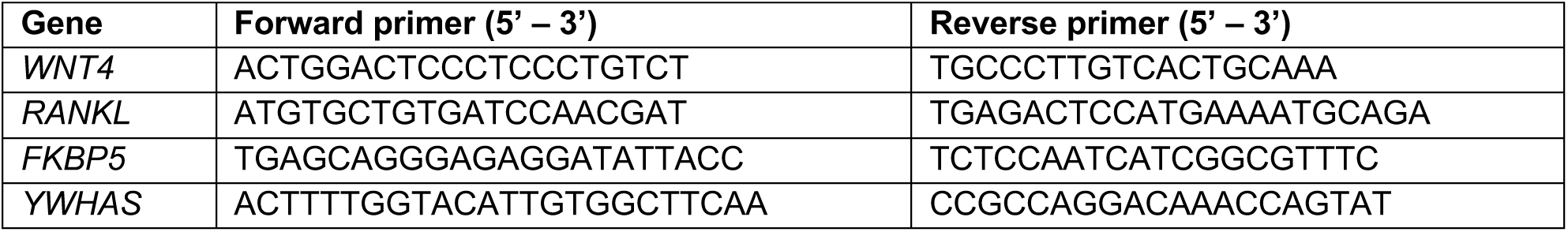
qRT-PCR primers.

### Western Blot

For Western Blot analysis, the cells were plated and treated in 6-well plates and lysed using 100μl lysis buffer (20µM Tris pH 8.0, 2µM EDTA pH 8.0, 0.5% NP40, 25µM sodium B-glycerophosphate, 100µM Sodium fluoride, 10mM sodium pyrophosphate). Protein concentrations were measured using the Pierce BCA protein Assay kit (Thermo Fisher, #23225) and a total of 30µg of protein for each sample was loaded on a 10% SDS-PAGE gel. Proteins were transferred on a 0.2 µm nitrocellulose membrane using the trans-blot turbo transfer system (Bio-Rad) and blocked with 1:1 diluted TBS Odyssey Blocking buffer (LI-COR Biosciences, #927-50100). Primary antibody directed against PR (1:1000, Thermo Fisher #MA5-16393, recognizing both PR-A and PR-B) and Actin (1:1000, MP biomedicals #08691001) were diluted in blocking buffer supplemented with 0.1% Tween-20. Primary antibody staining was performed O/N at 4°C followed by incubation with secondary antibodies (1:20000 IRDye 680L, #926– 6802 or 1:20000 IRDye 800CW LI-COR, #926–32211), in TBS supplemented with 0.1% Tween-20 for 1 hour at RT and detection was performed at 700nm and 800nm using an Odyssey Fc (LI-COR Biosciences).

### Generation of MCF7-PR and MCF7-PR-2/4/6xPRE-GFP lines

For lentivirus production, 5×10^6^ HEK293TN cells were plated in 10cm plates. The next day, the cells were transfected with 3µg packaging vector pCMVDR8.2 (Table 2, #10), 3µg RSV-rev (Table 2, #11), 3μg VSVg (Table 2, #12), and 8µg custom generated lentiviral CMV-PR or 2x/4x/6xPRE-GFP plasmids using PEI (Polysciences, #23966). Medium was refreshed after 24 hours, virus was collected after 48 hours, filtered through a 45µm filter, and diluted 1:4. MCF7 cells were infected in the presence of polybrene (1:2000, Merck Millipore #TR-1003-G). After 24 hours incubation, cells were, split, and selected for up to two weeks with blasticidin (10µg/ml Thermo Fisher, #11583677) or G418 (700µg/ml Gibco, #11811-031).

### Fluorescence-activated cell sorting (FACS)

For FACS experiments, MCF7-PR-2/4/6xPRE-GFP cells were plated in 6-well plates for analysis, or 10cm plates for sorting, in fresh DMEM supplemented with 5% stripped FBS, 48 hours prior to analysis or sorting. At 24 hours, the cells were treated with EtOH, 20nM R5020 (sorting) or a R5020 concentration range from 10pM to 100nM (analysis) in fresh DMEM medium supplemented with 5% stripped FBS. 24 hours after treatment, the cells were trypsinized, and pelleted. The cells were stained with 1µg/ml DAPI (Invitrogen, #D1306) in HF (2% Charcoal Stripped FBS in HBSS), washed and again resuspended in HF and then filtered through a 70µm filter. Sorting and analysis were performed on a FACSAria^TM^ III (BD, Franklin Lakes, NJ). For FACS sorting, the cells with intermediate GFP expression levels were sorted in 24-wells plates containing full medium +1% penicillin/streptomycin and 0.025M HEPES. Analysis of FACS results was performed using Flowjo (10.8.2). Gates for GFP positive cells were set using MCF7 wildtyoe and EtOH treated samples of 2xPRE-GFP. The gating strategy was expanded to all other samples.

### Imaging of PRE-GFP lines

For imaging, MCF7-PR-2/4/6xPRE-GFP cells were seeded on an 8-well chamber slide with glass bottom (Ibidi, #80827-90) containing DMEM medium supplemented with 5% stripped FBS. After 6 hours, medium was replaced with phenol red free DMEM supplemented with 5% charcoal stripped FBS, containing either EtOH or 20nM R5020. The next morning, phenol red free DMEM supplemented with 5% charcoal stripped FBS, 0.025M HEPES (Thermo Fisher, #11560496), and 500nM SiR-DNA was added and 4 hours later, cells were imaged at 37°C on an SP8 confocal microscope (Leica Microsystems) using a 25x water objective, 488 (8% laser power) and 633 (8% laser power) lasers, using a HyD detector (100 gain) for fluorescent signal with a 496-559 bandpass for GFP and a PMT3 detector (700 gain) for fluorescent signal with a 642-693 bandpass for SiR-DNA. Images were processed using Fiji [50].

### Statistical analysis

For quantification of PR nuclear abundance (Fig1), P-values were calculated using a one-way ANOVA followed by a Tukey’s multiple comparison test. For luciferase assays and qRT-PCR analyses, P-values were calculated using a 2-way ANOVA followed by a Tukey’s multiple comparison test, except for the analysis of *PGR* expression (SupFig1) where a one-way ANOVA followed by a Tukey’s multiple comparison test was used. All statistical analyses were performed and plotted using GraphPad Prism (10.0.0).

## Statements and declarations

### Acknowledgments

We thank Gonzalo Congrains Sotomayor (Swammerdam Institute for Life Sciences, University of Amsterdam) for providing support for our FACS experiments and Prof. Dr. Pernette Verschure (Swammerdam Institute for Life Sciences, University of Amsterdam) and Dr. Stieneke van den Brink (Hubrecht Institute, Utrecht) for kindly providing the MCF7 and T47D cell lines, respectively. We also thank Dr. Harm Krugers (Swammerdam Institute for Life Sciences) for sharing the Aldosterone and Dr. Bart van der Burg for his input and suggestions during the luciferase assay optimization. We thank our colleagues from the Developmental, Stem Cell & Cancer Biology (DSCCB) group for discussions and feedback during the project. RvA acknowledges funding support from the Netherlands Organization for Scientific Research (OCENW.KLEIN.169).

### Author contributions

M.T.A, A.L.v.B and R.v.A conceived the study. M.T.A. and M.W. designed and performed 2-12xPRE luciferase experiments, medium optimization experiments, and luciferase experiments with the mutated PRE consensus sequences in MCF7 cells.

T.v.d.W and M.T.A. designed and performed immunofluorescence and imaging experiments. M.T.A. designed and performed all other experiments. M.T.A. wrote the manuscript, with input from A.L.v.B and R.v.A. All authors approved submission.

## Notes

### Competing Interest Statement

The authors have declared no competing interest.

